# Multiple pathways to homothallism in closely related yeast lineages in the Basidiomycota

**DOI:** 10.1101/2020.09.30.320192

**Authors:** Alexandra Cabrita, Márcia David-Palma, Patrícia H. Brito, Joseph Heitman, Marco A. Coelho, Paula Gonçalves

**Affiliations:** Applied Molecular Biosciences Unit-UCIBIO, Departamento de Ciências da Vida, Faculdade de Ciências e Tecnologia, Universidade Nova de Lisboa, 2829-516 Caparica, Portugal; Department of Molecular Genetics and Microbiology, Duke University, Duke University Medical Center, Durham, NC 27710, USA.

## Abstract

Sexual reproduction in fungi relies on proteins with well-known functions encoded by the mating-type (*MAT*) loci. In the Basidiomycota, *MAT* loci are often bipartite, with the *P/R* locus encoding pheromone precursors and pheromone receptors and the *HD* locus encoding heterodimerizing homeodomain transcription factors (Hd1/Hd2). The interplay between different alleles of these genes within a single species usually generates at least two compatible mating types. However, a minority of species are homothallic, reproducing sexually without an obligate need for a compatible partner. Here we examine the organization and function of the *MAT* loci of *Cystofilobasidium capitatum*, a species in the order Cystofilobasidiales, which is unusually rich in homothallic species. We determined *MAT* gene content and organization in *C. capitatum* and found that it resembles a mating type of the closely related heterothallic species *Cystofilobasidium ferigula*. To explain the homothallic sexual reproduction observed in *C. capitatum* we examined HD-protein interactions in the two *Cystofilobasidium* species and determined *C. capitatum MAT* gene expression both in a natural setting and upon heterologous expression in *Phaffia rhodozyma*, a homothallic species belonging to a clade sister to *Cystofilobasidium*. We conclude that the molecular basis for homothallism in *C. capitatum* appears to be distinct from that previously established for *P. rhodozyma*. Unlike the latter species, homothallism in *C. capitatum* may involve constitutive activation or dispensability of the pheromone receptor and the functional replacement of the usual Hd1/Hd2 heterodimer by an Hd2 homodimer. Overall, our results suggest that homothallism evolved multiple times within the Cystofilobasidiales.

**Importance:** Sexual reproduction is important for the biology of eukaryotes because it strongly impacts the dynamics of genetic variation. In fungi, although sexual reproduction is usually associated with the fusion between cells belonging to different individuals (heterothallism), sometimes a single individual is capable of completing the sexual cycle alone (homothallism). Homothallic species are unusually common in a fungal lineage named Cystofilobasidiales. Here we studied the genetic bases of homothallism in one species in this lineage, *Cystofilobasidium capitatum*, and found it to be different in several aspects when compared to another homothallic species, *Phaffia rhodozyma*, belonging to the genus most closely related to *Cystofilobasidium*. Our results strongly suggest that homothallism evolved independently in *Phaffia* and *Cystofilobasidium*, lending support to the idea that transitions between heterothallism and homothallism are not as infrequent as previously thought. Our work also helps to establish the Cystofilobasidiales as a model lineage in which to study these transitions.

## Introduction

In Fungi, as in all eukaryotes, sexual reproduction is widespread, and some of the underlying mechanisms and molecular components are conserved even among distant lineages. The specific molecular pathways involved and the recognition systems responsible for triggering sexual reproduction may nonetheless vary greatly (1). Generally, sexual reproduction occurs through mating of two haploid individuals of the same species, possessing distinct mating types (2, 3). Mating types are defined by specific regions of the genome, the mating type (*MAT*) loci, which encode proteins responsible for triggering the major pathways leading to sexual development. Distinct mating types differ in the genetic content of the *MAT* loci. In basidiomycetes, mating-type determination relies on two *MAT* loci (named *P/R* and *HD*) that encode two different classes of proteins (4–6). The *HD* locus contains two divergently transcribed genes encoding homeodomain transcription factors and the *P/R* locus comprises pheromone receptor and pheromone precursor encoding genes - *STE3* and *MFA*, respectively (4, 5, 7). In the Ascomycota, only one *MAT* locus encoding transcription factors is required to determine mating-type identity. Hence, the participation of receptors and pheromones in the determination of mating type is found only in the Basidiomycota.

The *HD* and *P/R MAT* loci in Basidiomycota can be either genetically linked or unlinked in the genome of a given species. If these loci are unlinked, they may segregate independently during meiosis, leading to the generation of four distinct mating types among the haploid progeny, which is the hallmark of the tetrapolar breeding system (4, 5, 7). The bipolar breeding system results from linkage of the two *MAT* loci and yields only two mating types in the haploid progeny (4, 5, 7). Bipolar mating also takes place if one of the two (unlinked) *MAT* loci loses its function in determining mating-type identity, which has been reported occasionally for the *P/R* locus (4, 5, 7).

Although heterothallism involving the fusion between compatible mating types as a requirement for sexual reproduction is common, in some fungal species individuals are universally compatible, meaning that they can undergo sexual reproduction with any other individual, or even with or by itself, a pattern termed homothallism (3, 7, 8). In basidiomycetes few homothallic organisms have been found and even fewer have had the genetic basis of their homothallism elucidated (1, 3, 7, 8). Two relevant examples of basidiomycete yeasts with distinct molecular strategies that result in a homothallic sexual mode of reproduction are the human pathogenic yeast *Cryptococcus deneoformans* (9, 10) and *Phaffia rhodozyma* (11). *Cryptococcus deneoformans* exhibits a particular form of homothallism named unisexual reproduction, where cells of the same mating type can either fuse or undergo endoreplication of the entire genome resulting in a diploid nucleus that subsequently develops into hyphae, basidia, and gives rise to four haploid spores that are products of a meiotic event. Interestingly, the key genes for heterothallic reproduction like *MFA, STE3* and even *HD1* (*SXI1)* and *HD2* (*SXI2*) appear to be dispensable for unisexual reproduction of some *C. deneoformans* strains (10, 12, 13).

More recently, the genetic basis of homothallism was dissected in *Phaffia rhodozyma* in our laboratory (11). This astaxanthin-producing basidiomycetous yeast (14) belongs to the order Cystofilobasidiales, a lineage with an unusually high proportion of homothallic species (15–20), but comprising also heterothallic species and others for which sexual reproduction has not yet been observed. In *P. rhodozyma*, deletion mutants were used to show that the two pairs of *STE3* and *MFA* genes, and the single *HD1/HD2* gene pair present in the genome, are all required for robust sexual reproduction (11). The two pairs of pheromone and pheromone receptors turned out to have reciprocal compatibility, a single compatible *STE3* and *MFA* pair being sufficient for sexual reproduction. This is what might be expected if the *P/R* loci of two putative mating types were present in the same genome, in accord with the definition of primary homothallism. The only pair of Hd1 and Hd2 proteins encoded in the genome is also essential for sexual development, but the mode of action is not fully understood (11). This is because Hd1 and Hd2 are usually expected to heterodimerize to form functional transcription factors, but proteins encoded in the same *HD* locus do not normally form dimers. This imposes heterozygosity at the *HD* locus, with dimerization occurring only between proteins encoded by different gene pairs, as a condition for sexual development. In line with this, the Hd1 and Hd2 proteins of *P. rhodozyma* that are encoded in the same gene pair, do not appear to interact strongly, leading to the tentative conclusion that a weak interaction between the two proteins might suffice for function (11). Therefore, the *HD* locus configuration in *P. rhodozyma* is not fully aligned with the concept of primary homothallism, where the presence of two distinct pairs of *HD* genes supporting strong cross dimerization between Hd1 and Hd2 proteins would be expected.

Here, we examine in detail the content and function of the *MAT* loci of *Cystofilobasidium capitatum*, a homothallic species closely related to *P. rhodozyma*, in order to understand if there are common features between the molecular basis of homothallism in both species. We aim to shed some light on the diversity of molecular mechanisms through which homothallism can occur in the phylum Basidiomycota and improve the understanding of the evolution of mating patterns in the Cystofilobasidiales.

### RESULTS

### *MAT* loci in *Cystofilobasidium capitatum* and *Cystofilobasidium ferigula*

*Cystofilobasidium capitatum* and *Cystofilobasidium ferigula* belong to the order Cystofilobasidiales that also contains the genus *Phaffia*, for which two new species were recently described (21), in addition to six other genera (22). The phylogenetic relationships within the order were only recently clarified by a comprehensive genome-based phylogeny (21). This order contains a strikingly large number of homothallic species (approximately one third of those described so far; **Table S1 (10.6084/m9.figshare.13176422)**), a sexual mode uncommon in the Basidiomycota (3). In particular, *Phaffia* is composed entirely of homothallic species (21) while *Cystofilobasidium* comprises species with a variety of sexual properties, with *C. capitatum* and *C. intermedium* (23) representing the only two fully homothallic species in the genus. The remaining species of *Cystofilobasidium* are heterothallic except for *C. macerans*, which comprises strains exhibiting diverse sexual patterns (heterothallic, homothallic, and asexual) and *C. alribaticum* for which no sexual reproduction has been observed under the conditions tested (23). The genome-based phylogeny published for the Cystofilobasidiales (21) robustly supported *Phaffia* and *Cystofilobasidium* as sister genera, a relationship that is recapitulated in the phylogeny shown in **Fig. 1**, where a more limited number of species were included.

**Fig. 1.**
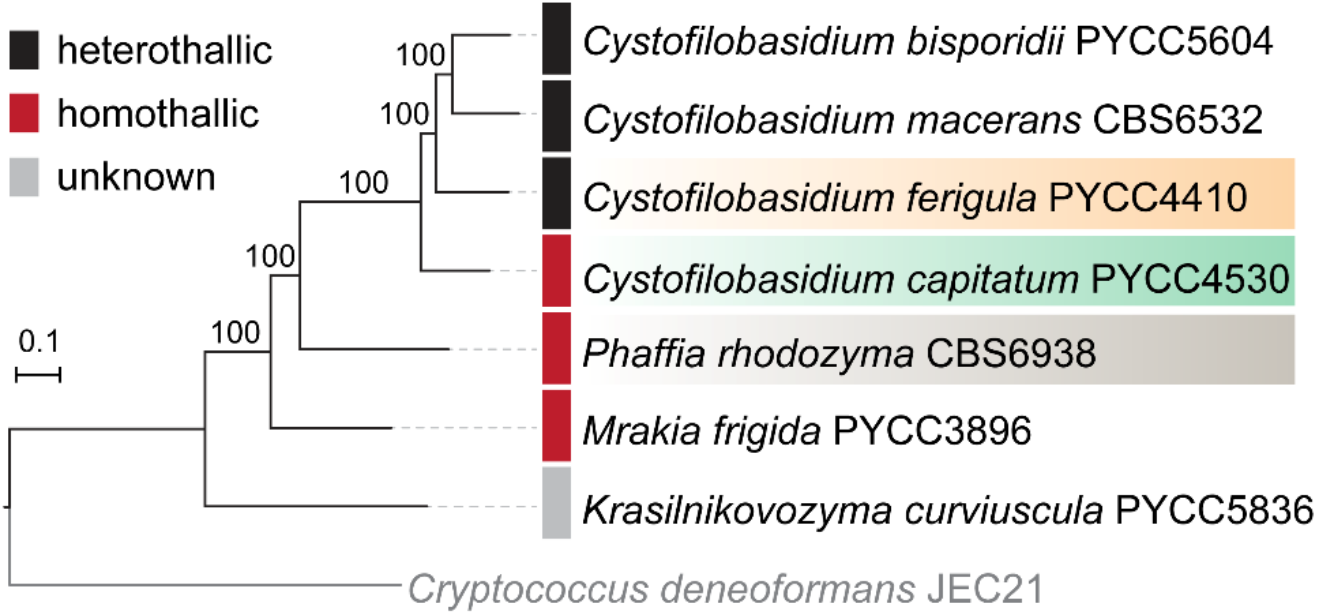
Maximum likelihood phylogeny reconstructed from the concatenated protein alignments of 1147 single-copy genes shared across the studied taxa and the outgroup represented by *Cryptococcus deneoformans*. Branch support was assessed by 1000 replicates of ultrafast bootstrap approximation (UFBoot) with NNI optimization, and branch lengths are given in number of substitutions per site.

The availability of draft genomes for several *Cystofilobasidium* species allowed us to examine the *MAT* loci of *C. capitatum* as well as that of the heterothallic species *C. ferigula* (**Fig. 2** and **Fig. S1**) and to compare these with *MAT* loci in the homothallic *P. rhodozyma* concerning gene content and organization (24). In the genome of *C. capitatum* PYCC 4530 a single pair of divergently transcribed homeodomain transcription factors (*HD1* and *HD2*) was found, as well as a single pheromone receptor gene (*STE3*) flanked by two identical pheromone precursor genes (*MFA1a* and *MFA1b*; **Fig 2**). These two sets of genes are located on different scaffolds and encode proteins with similar lengths to their counterparts in *P. rhodozyma* and the receptor is predicted to possess seven transmembrane domains as expected (**Fig. 2B and Fig. S2**). Both predicted Hd proteins have homeobox domains and nuclear localization signals (NLS; **Fig**.**S2**). However, the *MFA* genes in *C. capitatum* encode a 41 amino acid pheromone precursor protein where no site for N-terminal processing could be identified (**Fig. 2C**), which may compromise the formation of a mature pheromone (25). Furthermore, analysis of gene synteny conservation between *Phaffia* and *C. capitatum* indicates that the *P/R* locus of *C. capitatum* may be restricted to the region containing the pheromones/receptors (highlighted in yellow in **Fig. 2B**) and the *HD* locus most likely includes only the *HD1* and *HD2* genes, as observed in many other basidiomycetes (11, 26, 27).

**Fig. 2.**
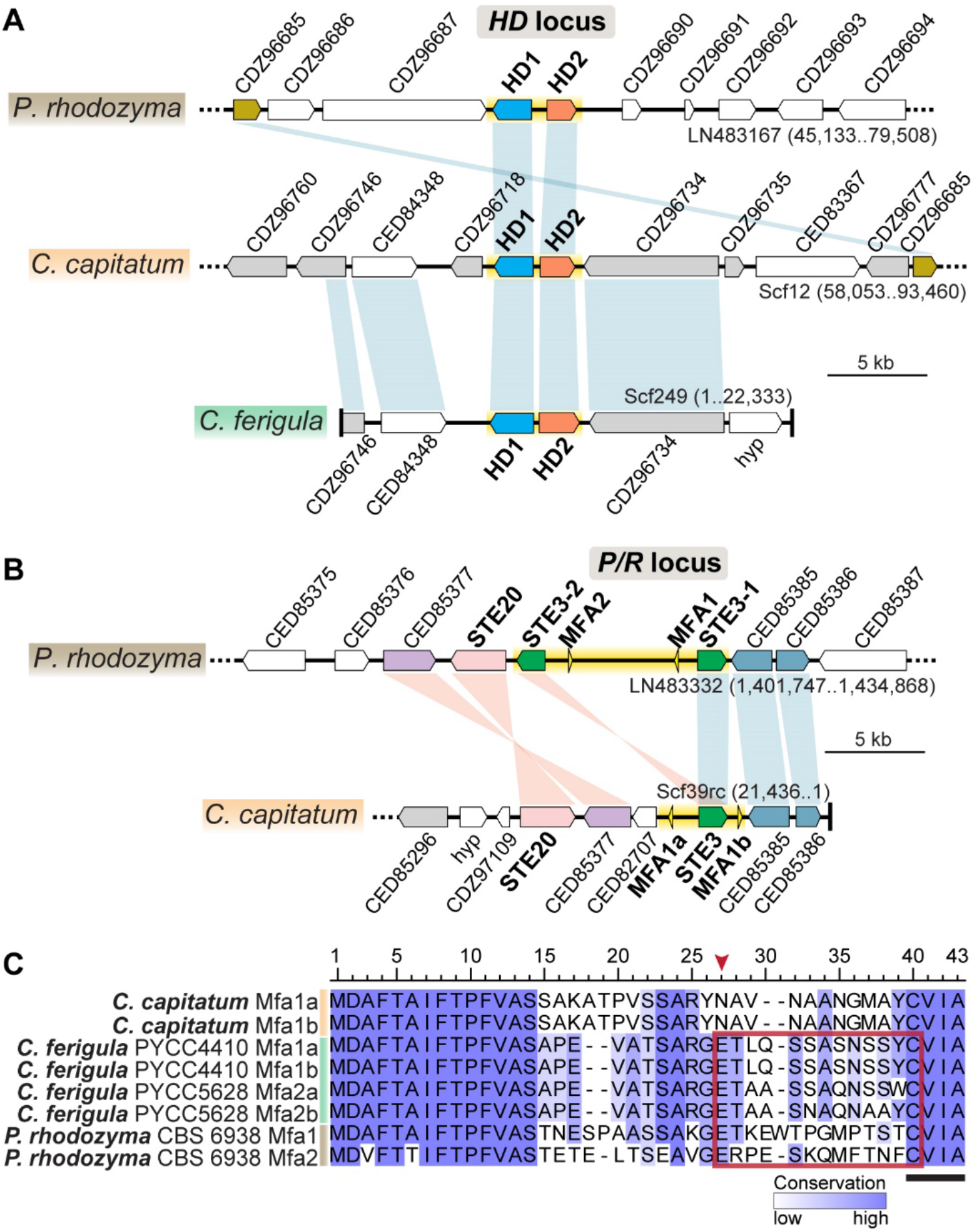
Gene content and organization of the (A) *HD* and (B) *P/R* mating-type loci in *C. capitatum* and *C. ferigula* (PYCC 4410), compared to *P. rhodozyma*. Genes are depicted as arrows denoting the direction of transcription. The genomic regions corresponding to the putative *MAT* loci are highlighted in yellow. Vertical bars connect orthologs that are in the same (blue) or inverted (pink) orientation. Non-syntenic genes are shown in white, and genes in grey in *C. capitatum* and *C. ferigula* are found scattered in the corresponding *P/R*- or *HD*-containing scaffold in *P. rhodozyma*, suggesting low level of synteny conservation beyond the core *MAT* regions. Gene names are based on top BLASTp hits in the *P. rhodozyma* genome and “hyp” denotes hypothetical proteins. The *P/R* locus of *C. ferigula* was omitted because the available genome assemblies are too fragmented in these regions to allow for a comparison. (C) Sequence alignment of pheromone precursors with amino acid positions colored in a blue gradient according to conservation. Sequences proposed as the putative mature pheromones are outlined by a red box, while those resembling the CAAX motif required for farnesylation are underlined. In *C. capitatum* the absence of a conserved position for N-terminal processing (the two charged amino acids indicated by a red arrowhead; ref 25), precludes the prediction of a mature pheromone sequence.

For *C. ferigula*, the genomes of two compatible mating types were obtained (PYCC 4410 and PYCC 5628). Findings concerning *MAT* gene content were similar to *C. capitatum* (**Fig. 2** and **Fig. S1**), and in line with the genetic composition typically found in haploid mating types of basidiomycetes. In *C. ferigula*, two genes encoding pheromone precursors were also found in each of the mating types, but in strain PYCC 5628 the two genes seem to encode slightly different mature pheromones. Because these genes are found in very small contigs in the current genome assemblies, it was not possible to determine their position in relation to the *STE3* gene nor the exact number of copies in the genome. It is conceivable that additional pheromone genes may become apparent when more complete assemblies of *C. capitatum* and *C. ferigula* genomes are available. As in *C. capitatum*, the *HD1/HD2* and the *STE3* genes were also found on different scaffolds in both *C. ferigula* strains, but the higher level of fragmentation of the current assemblies precludes a precise determination of the length and configuration of the *P/R* and *HD MAT* loci.

While heterothallic species are expected to harbour at least two different mating types with distinct alleles of *MAT* genes, this does not necessarily apply to homothallic species because there is no requirement for an operational self/nonself-recognition system in this case. To assess allele diversity of *MAT* genes in the *Cystofilobasidium* species under study, we obtained *MAT* gene sequences for as many strains as possible for both species. For *C. ferigula* two clearly distinct *STE3* alleles (sharing ∼51% amino acid sequence identity) could be recognized among the five strains examined, as expected for a heterothallic species (**Fig. 3A**). For *C. capitatum*, although several different alleles could be identified in the 11 strains examined, they were much less divergent (sharing ∼98% amino acid sequence identity) than *STE3* alleles known to encode proteins with different specificities, like those of *P. rhodozyma* (∼50% sequence identity; (11)) and *C. ferigula* (**Fig. 3A**).

**Fig. 3.**
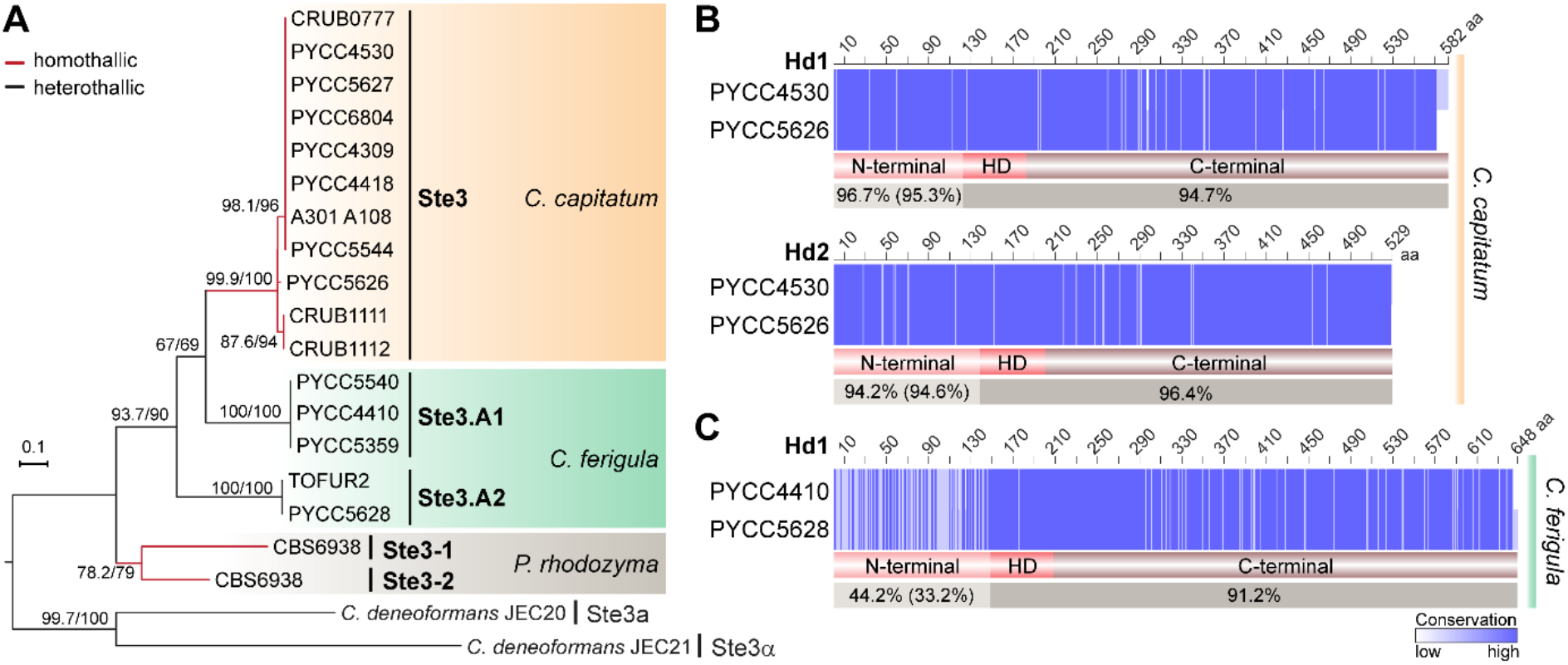
Sequence diversity of *STE3* and *HD* mating-type genes in *C. capitatum* and *C. ferigula*. (A) Maximum likelihood phylogeny of pheromone receptors (Ste3) obtained from various strains of *C. capitatum* (homothallic) and *C. ferigula* (heterothallic), along with the previously characterized Ste3 sequences of *P. rhodozyma* (homothallic) and *C. deneoformans* (heterothallic). The tree was inferred with the LG+F+G4 model of amino acid substitution and was rooted in the midpoint. Branch support values separated by a slash were assessed by 10,000 replicates of both the Shimodaira-Hasegawa approximate likelihood ratio test (SH-aLRT) and the ultrafast bootstrap approximation (UFBoot). Compared to the other species, the low sequence divergence of Ste3 sequences among *C. capitatum* strains suggest the absence of functionally distinct, mating-type specific, alleles in this species. (B) and (C) Sequence alignments of the *HD1* and *HD2* gene products of *C. capitatum* and *C. ferigula*. Sequence identity between each pair of Hd1 and Hd2 proteins is given for the variable (N-termini) and conserved (homeodomain and C-termini) regions with amino acid positions colored in a blue gradient according to conservation. The comparison between Hd2 proteins of *C. ferigula* was not performed because the *HD2* gene of PYCC 5628 is fragmented in the current genome assembly. Values in brackets for the N-terminal regions are the average identity as calculated from different allele products (see **Fig. S3C** for details on number of alleles). In *C. capitatum*, variable amino acid positions are evenly distributed throughout the length of the Hd1 and Hd2 proteins. In contrast, the N-terminal region of Hd1 in *C. ferigula* is comparatively more variable, as commonly observed in Hd1 and Hd2 proteins of other heterothallic basidiomycetes (5, 7, 12, 26).

The comparison of *C. capitatum* Hd proteins suggest the existence of three main Hd variants in the species with a degree of divergence between them that is lower than observed for functionally different alleles of heterothallic species (**Fig. S3B and S3C)**. Moreover, for the two proteins that could be examined over their entire length, the differences in the sequences are distributed homogeneously throughout the protein (**Fig. 3B**), like observed for *P. rhodozyma* (11). This is unlike the divergence observed in Hd1 sequences of *C. ferigula* and of other heterothallic basidiomycetes, which is more extensive and concentrated in the N-terminal region responsible for self/nonself-recognition (**Fig. 3C**).

*C. ferigula* was reported to be bipolar, as only two different mating types had been identified so far (16). This would mean that each *STE3* allele is expected to be linked to a single *HD* allele, defining two mating types. However, as shown in **Fig. S3C**, our analysis uncovered three *HD* alleles, instead of the expected two, and the *HD*.*B1* allele appears associated with the two receptors in different strains, whereas the *STE3*.*A1* allele is associated with *HD*.*B1* and *HD*.*B3*. From these observations it seems more likely that *C. ferigula* may have a tetrapolar mating system, in contrast to previous assumptions (16).

### Involvement of the Hd proteins of *C. capitatum* in the homothallic sexual cycle

In heterothallic basidiomycetes, the Hd1 and Hd2 proteins encoded in the *HD* locus control the later stages of sexual reproduction through heterodimerization of non-allelic Hd1 and Hd2 proteins brought together by cell fusion (4, 12, 26). Previous studies in our laboratory concerning the molecular mechanisms of sexual reproduction of *P. rhodozyma* revealed that both the Hd1 and Hd2 proteins are required for normal sporulation (which for this species consists of basidia formation), and likely act through heterodimerization despite the weak nature of their interaction (11).

To understand if *C. capitatum* resembled *P. rhodozyma* in this respect, the yeast two-hybrid assay was employed to assess the ability of the Hd1 and Hd2 proteins of *C. capitatum* to interact with each other. *HD1* and *HD2* cDNAs were isolated from *C. capitatum* strain PYCC 5626 and utilized for the construction of the Gal4 fusion genes for this assay. The results of the assay, presented in **Fig. 4**, are consistent with a complete absence of interaction between the Hd1 and Hd2 proteins of *C. capitatum*, unlike the results for *P. rhodozyma* (11). Notably, a strong homodimerization was detected for the Hd2 protein from this species. (**Fig. 4**). Weak homodimerizations of Hd proteins were also previously observed for *P. rhodozyma* (11). For *C. ferigula* a strong heterodimerization of the Hd1 and Hd2 proteins derived from strains of different mating types (PYCC 5628 and PYCC 4410, respectively) was noted (**Fig. 4**), in line with observations in other heterothallic basidiomycete species (12, 26). Consistently, β-galactosidase activity resulting from activation of the *lacZ* reporter gene was similar to the positive control both for heterodimerization of Hd proteins from *C. ferigula* and for homodimerization of Hd2 in *C. capitatum* (**Fig. 4B**). Homodimerization of Hd1 in *C. capitatum* could not be tested, because the construct of the fusion protein between Hd1 and the Gal4 DNA binding domain could not be stably expressed in the pertinent *S. cerevisiae* strain.

**Fig. 4.**
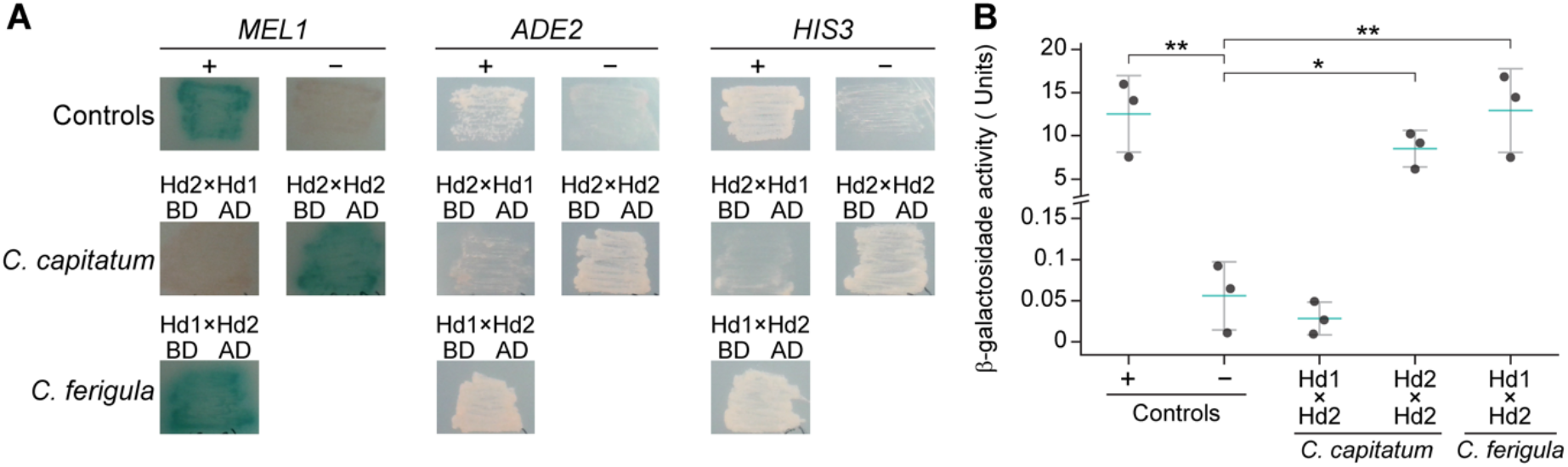
Results of the Yeast Two-Hybrid assay for Hd proteins of *C. capitatum* and *C. ferigula*. (A) Qualitative results concerning the dimerization of Hd1 and Hd2 of *C. capitatum* and of *C. ferigula*, as well as homodimerization of Hd2 of *C. capitatum*, through the activation of the reporter genes *MEL1, ADE2*, and *HIS3*. Activity of the three reporter genes was tested separately in appropriate media, namely containing X-α-Gal and all required supplements *for MEL1*,or lacking supplementation with adenine (*ADE2)* or histidine *(HIS3)*. Activity of the two latter reporter genes is denoted by the ability of the strains to grow on these media, while *MEL1* expression results in the formation of blue colonies. In *C. ferigula* the Hd1 protein sequence is derived from strain PYCC 5628 and the Hd2 protein is derived from strain PYCC 4410. (B) Quantitative results of the β-galactosidase assay of interactions shown in panel A. Each datapoint is the average of the activity measured in two replicate assays; the three datapoints shown for each strain represent β-galactosidase activities measured in reactions stopped after 2, 6 and 24 hours (see also Table S8 (10.6084/m9.figshare.13176422)). Blue bars denote the mean and vertical bars the standard error of the means. The interaction between a fusion protein containing the Gal4 activation domain fused to the SV40 large T-antigen and a fusion between Gal4 binding domain and p53 was used as positive control, while the negative control employed the same Gal4 activation domain fusion in combination with a fusion between Gal4 binding domain and lamin. Plasmids encoding the positive and negative control proteins were provided with the Matchmaker Gold Yeast Two-Hybrid System, by Takara Bio USA. AD, activation domain of Gal4; BD, DNA binding domain of Gal4. Significant differences between means were calculated using the Tukey’s HSD test (*p<0.05, **p<0.01).

### The *HD* locus of *C. capitatum* partially complements a *P. rhodozyma HD* deletion mutant

The results obtained in the yeast two-hybrid assay suggest that homodimerization of Hd2 may play a role in homothallic sexual development in *C. capitatum*, possibly functionally replacing the usual Hd1/Hd2 heterodimer. To investigate this, we set out to assess function of the *C. capitatum HD* locus by heterologous expression in a *P. rhodozyma HDΔ* mutant.

Integration of the complete *HD* locus of *C. capitatum* strain PYCC 4530 in the rDNA locus of the *P. rhodozyma HDΔ* mutant (construct *HDΔ+HD1/HD2-PYCC4530*) resulted in a very weak but consistent recovery of sporulation (**Table 1, Table S2 and Table S3 (10**.**6084/m9**.**figshare**.**13176422)**). Because no interaction between the Hd1 and Hd2 proteins could be detected in the yeast two-hybrid assay but instead strong homodimerization of the Hd2 protein was observed, we subsequently decided to assess whether expression of *HD2* alone was sufficient to restore sexual development of the *P. rhodozyma HDΔ* mutant. To this end, a construct containing in addition to the intergenic region only a residual (344 bp) 5’portion of the *HD1* gene (excluding the homeodomain; *HDΔ+HD2-PYCC4530*) was used to transform the *P. rhodozyma HDΔ* mutant. In this mutant that expresses only the Hd2 protein, sporulation levels like those observed upon transformation of the complete *C. capitatum HD* locus were observed (**Table 1, Table S2 and Table S3 (10**.**6084/m9**.**figshare**.**13176422)**), suggesting that only the Hd2 protein is required for the observed complementation.

**Table 1.**
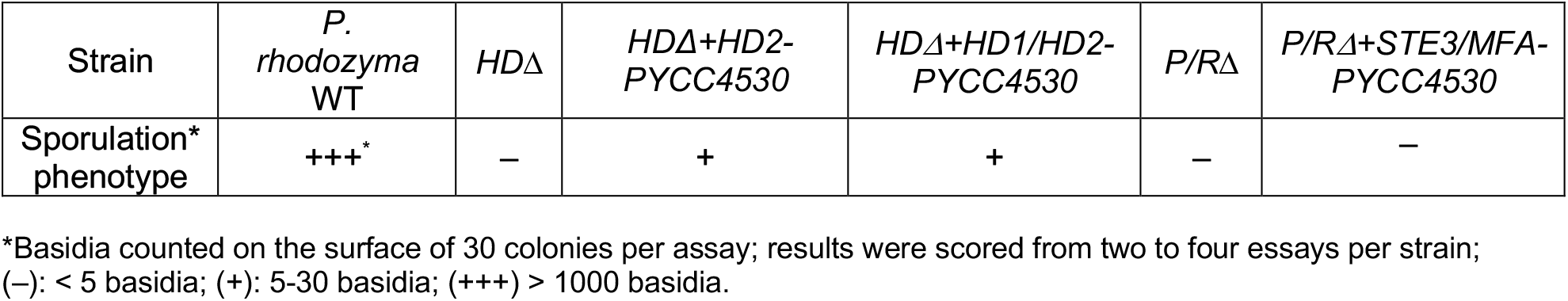
Complementation of *P. rhodozyma MAT* loci mutant strains with *C. capitatum* genes. Sporulation (basidia formation) patterns were scored qualitatively using the criteria explained as given in the key.

### The *HD1 and HD2* genes of *C. capitatum* are differently expressed during growth in sporulation conditions and in *P. rhodozyma* mutants

To substantiate the hypothesis that Hd2 might be the sole important component in the *HD* locus of *C. capitatum*, we compared the expression levels of *HD1* and *HD2* in *C. capitatum* PYCC 5626, in sporulation conditions, and in the *P. rhodozyma* mutant containing the complete *HD* locus of *C. capitatum* PYCC 4530 (construct *HDΔ+HD1/HD2-PYCC4530*). The results, depicted in **Fig. 5**, show that of the *HD* gene pair, only the *HD2* gene seems to be substantially expressed in the *C. capitatum* strain. Heterologous expression of *HD2* in the *P. rhodozyma* mutant, although much lower than in *C. capitatum*, can be easily detected, while heterologous expression of *HD1* seems to be only vestigial.

**Fig. 5.**
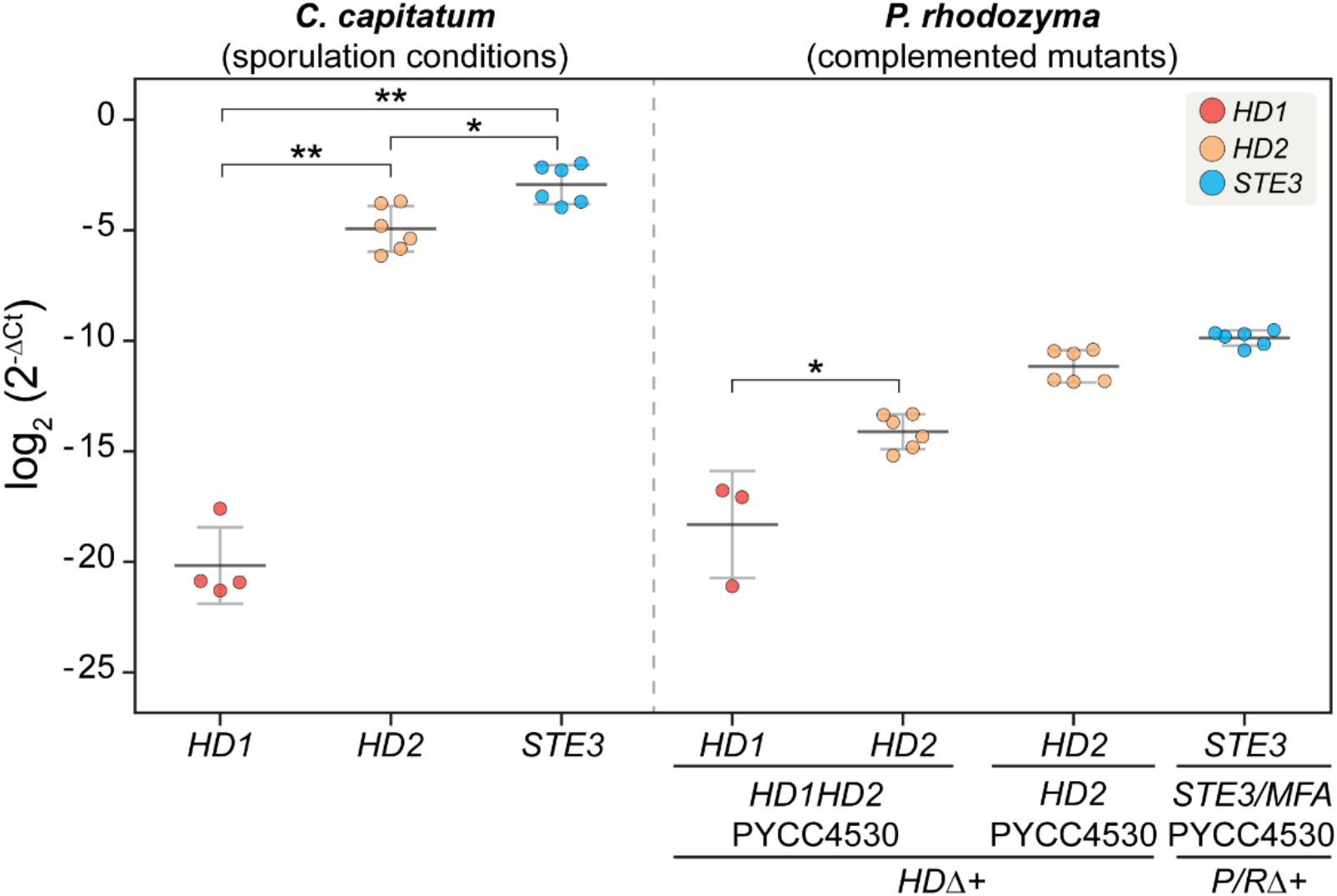
Real time quantitative PCR results of *MAT* gene expression in *C. capitatum* under sporulation conditions and in *P. rhodozyma* complemented mutants. Expression of *MAT* genes is given as log2 fold difference relative to the expression of actin in each strain. These results are derived from two biological replicates, each assayed in triplicate, resulting in six datapoints for each measurement. For *HD1* expression assays less than six dots are plotted because in some reactions *HD1* expression was undetectable. Significant differences between the expression of different genes were calculated with the Mann-Whitney test (*p<0.05, **p<0.01).

### *P/R* locus of *C. capitatum* does not restore sporulation in a *P. rhodozyma* cognate deletion mutant

Because Hd function could be assessed using heterologous expression, a *P. rhodozyma* deletion mutant of both *P/R* clusters (*P/RΔ*) was similarly used as host for the *P/R* locus of *C. capitatum* strain PYCC 4530, encompassing the *STE3* and *MFA1*a genes and the respective native promotor regions (*P/RΔ+STE3/MFA-PYCC4530*). However, integration of the *P/R* locus of *C. capitatum* in the rDNA of the *P. rhodozyma P/RΔ* mutant failed to restore sporulation (**Table 1 and Table S2 (10**.**6084/m9**.**figshare**.**13176422)**), although the *C. capitatum STE3* gene was expressed in *P. rhodozyma* (**Fig. 5**). Interestingly, *STE3* is the most expressed among *MAT* genes of *C. capitatum* PYCC 5626 (**Fig. 5**), suggesting it may have a role in sexual reproduction despite our inability to observe it in the heterologous setting.

### DISCUSSION

The main aim of this study was to ascertain to which extent common features could be found between the molecular bases of homothallism in different species of the Cystofilobasidiales, a lineage in the Basidiomycota particularly rich in species exhibiting this uncommon sexual behaviour. The molecular basis of homothallism was previously dissected in the genetically tractable Cystofilobasidiales species *P. rhodozyma* (11). Here, we characterized the *MAT* loci of a second homothallic species belonging to a sister genus, *C. capitatum*, by examining the structure of the loci in the available genome of strain PYCC 4530 and by comparing *MAT* gene sequences in a total of 11 *C. capitatum* strains. The most striking difference between the *MAT* loci in *P. rhodozyma* and *C. capitatum* is the presence in the latter species of a single pheromone receptor gene, instead of the two distinct and functionally complementary sets of pheromone receptor and pheromone precursor genes found in *P. rhodozyma*. Therefore, the *C. capitatum MAT* gene content resembles a haploid mating type of a heterothallic species (4), while that of *P. rhodozyma* is reminiscent of a fusion between two mating types (11). However, like in *P. rhodozyma*, no evidence for functionally distinct variants (alleles) of *MAT* genes that might form different mating types were found in *C. capitatum*, consistent with its homothallic behaviour. We subsequently devised various experimental approaches to try to bring to light functional features of *MAT* genes in *C. capitatum*. In these experimental approaches two *C. capitatum* strains were used: in addition to sequenced strain PYCC 4530, strain PYCC 5626 was employed for the isolation of cDNAs because unlike other strains, it readily sporulates in different experimental conditions.

*P. rhodozyma* can be described as a primary homothallic basidiomycete, with reciprocal compatibility between the pheromones and receptors of the two *P/R* clusters and with a weak heterodimerization between the only pair of Hd1 and Hd2 proteins being responsible for triggering of its sexual cycle, with possible hints at homodimerization events (11). Therefore, in *P. rhodozyma*, the *P/R* system seems to work similarly to what might be expected for a heterothallic mating type, while the *HD* locus is apparently operating through weak dimerization between adjacently encoded Hd1 and Hd2. This dimerization normally occurs only between proteins encoded by different *HD* alleles. Hence, the weak interaction in *P. rhodozyma* may have evolved to support sexual development in the absence of a second *HD* allele. Because the *MAT* loci in *C. capitatum* have the same gene content as haploid mating types of heterothallic species, in this case both the *P/R* and the *HD* loci probably underwent changes in their mode of operation to permit homothallism. For the *HD* locus, the complete absence of interaction between the Hd1 and Hd2 proteins, the formation of Hd2 homodimers, complementation results of the *P. rhodozyma HDΔ* mutant and the very low expression of the *HD1* gene suggest that the *HD* locus is relevant for sporulation but also that regulatory functions normally fulfilled by the Hd1/Hd2 heterodimer may have been taken over by the Hd2 homodimer. Hd1 homodimerization was not tested due to technical difficulties. Although it would have been very interesting to be able to test also Hd1 homodimerization, this lost importance when gene expression experiments showed that *HD1* expression was barely detectable. Interestingly, the α2 transcription factor, which is the Hd1 *S. cerevisiae* homologue, is capable of forming both a homodimer that represses transcription of **a**-specific genes in haploid α-cells, and a heterodimer with the a1 transcription factor (Hd2 homologue) that represses haploid-specific genes in diploid cells (28). This suggests that these proteins can operate both as homodimers and as heterodimers. However, to our knowledge, functional Hd1 or Hd2 homodimers have not been reported so far in basidiomycetes, although there was some evidence for homodimerization of Hd proteins in *P. rhodozyma* (11). Our hypothesis for the mode of action of the *HD* locus predicts that Hd1 might be dispensable. In some heterothallic basidiomycetes, it has been shown that the Hd1 and Hd2 proteins hold different functional domains that are essential for function of the transcription regulation complex (29–31). One such species is the mushroom *Coprinopsis cinerea*, where it has been shown that Hd1 contributes with the NLS region, allowing the heterodimer to be transported into the nucleus, while the homeodomain of Hd2 is required for binding of the complex to DNA (29, 32). Hence, in this case, formation of a heterodimer is required for function. In *C. capitatum*, Hd2 possesses both an NLS and a homeodomain so in that perspective it seems possible that Hd2 may indeed play a role in the homothallic life cycle of *C. capitatum* that is sufficient and independent of Hd1. It is possible that this hypothesized function of Hd2 evolved recently in this species, which could explain the fact that the *HD1* is not pseudogenized. Alternatively, the Hd1 may have acquired a function unrelated to sexual reproduction, which might entail its expression under different conditions, and justify its maintenance in the genome. Despite multiple attempts, the methods used to transform *P. rhodozyma* failed to yield transformants of *C. capitatum*, which precluded the possibility of testing the hypothesis regarding the role of Hd2 in this species by deletion of the *HD2* gene.

Introduction of the *P/R* locus into the *P. rhodozyma P/RΔ* mutant failed to restore sporulation of the cognate mutant, even though expression of the *STE3* gene in the heterologous setting could be demonstrated. The explanations for this are presently unclear, since the ability of Ste3-like receptors to activate heterologously a sexual development pathway over a much larger phylogenetic distance was previously demonstrated (33). As to the mode of action of the receptor in its normal setting, the possibility that it could be constitutively active should be considered, in line with the fact that the predicted pheromone precursor genes lack the N-terminal processing signals, likely precluding the formation of an active pheromone. Mutations within the pheromone receptor that can lead to constitutively active receptors have been previously reported, resulting in a bypass of the pheromone for sexual reproduction (25, 33–36). Mutations in the pheromone that allow it to activate the receptor encoded in the same *P/R* cluster are also known (25, 37). In fact, mutations that lead to self-activation or self-compatibility have been proposed to form the basis for the transition from a tetrapolar to a bipolar mating system, or even from a heterothallic to a homothallic mating behaviour (3, 5) for some species. If the pheromone system does not operate normally in *C. capitatum*, this may be a reason for the lack of complementation of the *P. rhodozyma P/R*Δ mutant by the *C. capitatum P/R* locus.

Hence, taken together, our results suggest that *P. rhodozyma* and *C. capitatum* attained homothallic breeding systems through different mechanisms, which is consistent with the hypothesis that the *P. rhodozyma* mating system arose at the origin of the genus, possibly by fusion of *P/R* loci of compatible mating types and the adaptation of the HD dimerization system. *C. capitatum*, on the other hand, has an extant makeup of its *MAT* loci that suggests it may have evolved from a heterothallic mating type, in which the Hd2 homodimer acquired a prominent role. If a mature pheromone is formed despite the absence of an N-terminal processing signal, it might be that the pheromone-receptor pair in *C. capitatum* have become self-activating. On the other hand, if no mature pheromone is formed, the pheromone receptor gene might have bypassed the pheromone requirement and is now constitutively active. Whatever the case may be, the *P/R* system still likely has a role in some aspects of the sexual development, which in agreement with the fact that *STE3* gene seems to be highly expressed. Although we cannot discard the hypothesis that the receptor is fulfilling a role unrelated to sexual reproduction, and while such receptors have been reported (38), they were not associated with pheromone precursor genes.

The different particularities of homothallism in the two Cystofilobasidiales species studied so far are suggestive of remarkable levels of plasticity in the evolution of sexual reproduction in this order. It will be interesting to conduct similar studies in other homothallic species of this order, which would allow us to get a more complete insight in the array of mechanisms involved as well as possible genomic rearrangement that may have been involved in the transitions between heterothallic and homothallic species. Having uncovered *P. rhodozyma* as a viable host for heterologous expression, opens the possibility of assessing the functionality of other *MAT* proteins from uncharacterised species in this order.

### MATERIALS AND METHODS

### Strains and culture conditions

*P. rhodozyma, C. capitatum* and *C. ferigula* strains (**Table S4 (10**.**6084/m9**.**figshare**.**13176422)**) were routinely grown in YPD medium (2% Peptone, 1% Yeast Extract, 2% Glucose, 2% Agar) at 17 to 20°C. For the preparation of electrocompetent cells, *P. rhodozyma* strains were grown in YPD liquid medium at 20°C, and transformants were incubated at 17°C in selective medium, consisting of YPD plates supplemented with the appropriate antifungal drugs (100 μg/ml of geneticin and/or 100 μg/ml of zeocin and/or 100 μg/ml of hygromycin B).

*Escherichia coli* strain DH5α (Gibco-BRL, Carlsbad, CA, USA) was used as a cloning host and was grown at 37°C in LB medium (1% NaCl, 1% Tryptone, 0.5% Yeast Extract and 2% Agar for solid medium) supplemented with ampicillin at 100 μg/ml when appropriate.

### DNA extraction, genome sequencing and assembly

Genomic DNA of *C. ferigula* PYCC 5628 was extracted from single cell-derived cultures using the ZR Fungal/Bacterial DNA MiniPrep kit (ZYMO Research). DNA samples were quantified using Qubit 2.0. Genome sequencing was carried out by commercial providers, at the Genomics Unit of Instituto Gulbenkian de Ciência, and at the Sequencing and Genomic Technologies Core Facility of the Duke Center for Genomic and Computational Biology. Two short insert-size libraries (∼500 bp) were prepared with the Nextera Kit and subsequently sequenced using the Illumina MiSeq and HiSeq2500 systems to generate paired 300- and 151-nt reads, respectively. After adaptor clipping using Trimmomatic (v0.36) the two sets of reads were assembled with SPAdes (v3.11.1) (39) (with parameters: “--careful” to reduce the number of mismatches and short indels in the final assembly, and the k-mer sizes: 21, 33, 55, 77, 99, 127, automatically selected based on read length). Genome assembly quality was assessed by the QUAST (v.5.0.2) (40), and gene models were predicted ab initio using Augustus (41) trained on *Cryptococcus neoformans*. Genome sequencing data generated, and final genome assembly statistics are given in **Table S5 (10**.**6084/m9**.**figshare**.**13176422)**.

### Identification of mating-type genes and synteny analyses

Scaffolds containing *MAT* genes, namely the homeodomain transcription factors (*HD1*/*HD2*) and the mating pheromones (*MFA*) and receptors (*STE3*), were identified by BLASTP or TBLASTN in the genome assemblies of *C. capitatum* PYCC 4530 (=CBS 7420; BioProject: PRJNA371774) and *C. ferigula* PYCC 4410 (BioProject: PRJNA371786), and in the newly obtained assembly of *C. ferigula* PYCC 5628 (BioProject: PRJNA371793). Well-annotated *P. rhodozyma MAT* proteins (11) were used as search query. The retrieved *MAT* genes were manually reannotated if required and analysed further: (i) the transmembrane regions in the pheromone receptor protein were predicted by HMMTOP software (42); (ii) the Homeodomain regions in Hd1 and Hd2 proteins were predicted by InterPro server (43) and compared to the previously characterized homeodomain proteins in Pfam database; (iii) nuclear localization signals (NLS) and coiled-coil motifs were identified in the complete Hd1 and Hd2 sequences using, respectively, the SeqNLS (with a 0.8 cut-off) (44) and Jpred4 (45) (see **Figs. S1B** and **S1C** and **Fig. S2**). Synteny between *MAT* regions of different strains and species was based on bidirectional BLAST analyses of the corresponding predicted proteins. The short pheromone precursor genes in the genomes of *C. ferigula* and *C. capitatum* were identified manually as they usually fail automatic detection.

### Species and MAT gene phylogenies

A phylogenetic analysis representing major lineages within Cystofilobasidiales was inferred on a concatenated protein dataset of single copy core genes of four *Cystofilobasidium* species, *Phaffia rhodozyma* CBS 6938, *Mrakia frigida* PYCC 3896, *Krasilnikovozyma curviuscula* PYCC 5836, and the outgroup *Cryptococcus deneoformans* JEC21 (**Table S4** (10.6084/m9.figshare.13176422)). Orthologous clusters were inferred with all-against-all BLASTP (NCBI Blast-2.2) searches and the Markov cluster algorithm (OrthoMCL v1.4; (46)) with inflation factor (F) of 1.5, and minimum pairwise sequence alignment coverage of 50% implemented in GET_HOMOLOGUES package (47). Clusters present in single copy in all analyzed genomes were retained, aligned with MAFFT v7.407 using the G-INS-I method and default parameter values (48), trimmed with BMGE v1.12 using the amino acid option (49), and finally concatenated into a single data set. The species phylogeny was inferred with IQ-TREE v1.6.12 (50) using maximum-likelihood (ML) inference under a LG+F+I+G4 model of sequence evolution. ModelFinder (51) was used to determine best-fit model according to Bayesian Information Criterion (BIC) and branch support was estimated using ultrafast bootstrap approximation (UFBoot) with NNI optimization (52), both implemented in IQ-TREE package.

To analyse the *MAT* gene content across strains of *C. capitatum* and *C. ferigula*, protein sequences of the *HD1, HD2* and *STE3* genes were retrieved from the genome assemblies and aligned separately. Conserved regions were used to design primers to amplify the corresponding genomic regions across the available strains of each species (**Table S6 (10.6084/m9.figshare.13176422)**). These regions include a ∼ 870-bp region of the *STE3* gene, and a ∼1.5-kb fragment encompassing the 5′ end and intergenic regions of the *HD1* and *HD2* genes (**Table S6 (10.6084/m9.figshare.13176422) and Fig. S3**). Genomic DNA was extracted through a standard Phenol-Chloroform method (53) and the regions of interest were PCR-amplified, purified using Illustra GFX PCR DNA and Gel Band Purification Kit (GE Healthcare Life Sciences), and then sequenced by Sanger Sequencing, at STABVida (Portugal). For phylogenetic analysis of *MAT* genes, amino acid or nucleotide sequences were individually aligned with MAFFT v7.310 (48) using the L-INS-i strategy (--localpair --maxiterate 1000) and poorly aligned regions were trimmed with TrimAl (--gappyout) (54). The resulting alignments were input to IQ-TREE v.1.6.5 (50) ML phylogenies using best-fit models automatically determined by ModelFinder (51) (parameter: -m MFP). The exact model employed in tree reconstruction is given in the respective figure legends. Branch support values were obtained from 10,000 replicates of both UFBoot (52) and the nonparametric variant of the approximate likelihood ratio test (SH-aLRT) (55). In addition, the option “-bnni” was employed to minimize the risk of overestimating branch supports with UFBoot when in presence of severe model violations. The resulting phylogenies were midpoint rooted and graphically visualized with iTOL v5.5.1 (56).

### Yeast Two-Hybrid assay

To assess the interaction between the Hd1 and Hd2 proteins, the Matchmaker Gold Yeast Two-Hybrid System kit (Takara) was used. In this system, fusion proteins containing the Gal4 DNA binding domain (BD) are expressed from plasmid pGBKT7 in *MAT***a** haploid strain Y2HGold, while fusion proteins containing the Gal4 activation domain (AD) are expressed from plasmid pGADT7 in *MAT***α** haploid strain Y187. Diploid strains expressing both AD and BD fusion proteins are used to test interactions and are obtained by mating the haploid strains expressing each of the fusion proteins of interest. Hence, coding DNA sequences of the pertinent *HD* genes were cloned into pGADT7 and pGBKT7, so as to yield plasmids expressing the desired fusion proteins.

Synthetic genes designed using the coding DNA sequences of the *HD1* and *HD2* genes of *C. ferigula* and were synthesised at Eurofins Genomics (Germany). The *HD1* and *HD2* gene sequences of strains PYCC 4410 and PYCC 5628 from *C. ferigula* respectively were adapted to the *S. cerevisiae* codon usage (**Fig. S4**). The synthetic genes were obtained as inserts of pEX-A258 plasmids.

cDNAs of *HD1* and *HD2* from *C. capitatum* were obtained from total RNA isolated from strain PYCC 5626, briefly as follows. Strain PYCC 5626 was cultivated in GSA medium (0.2% Glucose, 0.2% Soytone) in 10% of the flask volume, for 8 days at 20°C and 90 rpm (Sartorius Certomat IS incubator), until inspection under the microscope revealed the presence of teliospores (16). RNA extraction was performed using the *ZR Fungal/Bacterial RNA MiniPrep kit* (by ZYMO Research), with a single step of in-column DNase I Digestion to free the RNA samples of genomic DNA. cDNA was synthesized from total mRNA using Maxima H Minus Reverse Transcriptase (by Thermo Scientific) and oligo (dT)20 as primer, and synthesis of the second DNA strand was performed using specific primers for the complete *HD1* and *HD2* genes (**Table S6 (10**.**6084/m9**.**figshare**.**13176422)**). The fragments corresponding to the *HD1* and *HD2* cDNA sequences of strain PYCC 5626 were sequenced by Sanger Sequencing, at STABVida (Portugal). The protein sequences of Hd1 and Hd2 of strain PYCC 5626 were aligned with those of strain PYCC 4530 (**Fig. 3**.**B**) using the software MUSCLE (implemented in the software Unipro UGENE v1.30.0 (57)) and the level of intraspecific variability was calculated.

The *HD1* and *HD2* complete synthetic genes from *C. ferigula* and cDNA fragments from *C. capitatum* were amplified using primers that contained 40 bp tails at their 5’ ends that correspond to the flanking regions of the Multiple Cloning Sites present in pGADT7 and in pGBKT7 (**Table S6 (10**.**6084/m9**.**figshare**.**13176422)**). Plasmid pGADT7 was then linearized at the Multiple Cloning Site by digestion with Cla I (Thermo Scientific), while pGBKT7 was linearized by digestion with Pst I (Thermo Scientific). *S. cerevisiae* strains Y187 and Y2HGold were transformed with inserts and linearized vectors using the transformation method described in the Yeastmaker Yeast Transformation System 2 protocol. Transformants were selected in appropriate media (Yeast Nitrogen Base without amino acids, with appropriate supplements (**Table S7 (10**.**6084/m9**.**figshare**.**13176422)**)) at 30°C. Transformantions with linearized pGBKT7 and the *C*.*capitatum* Hd1 insert resulted in numbers of transformants similar to those observed in other transformations. However, unlike in other transformations, all transformants investigated harboured plasmids without insert. This was observed only for this combination of vector and insert and persisted in different transformation attempts.

Mating of haploid *S. cerevisiae* strains was performed by incubating a single colony from each of the two haploid transformants to be mated in 200 μl of YPD medium at 30°C, for 24 hours, at 250 rpm. After incubation, the cells were recovered and thoroughly washed with distilled sterile water and plated on appropriate selective media (**Table S7 (10**.**6084/m9**.**figshare**.**13176422)**).

In the Matchmaker Gold Yeast Two-Hybrid System the *MEL1, ADE2 and HIS3* reporter genes are used for qualitative assessment of interactions using plate assays, while the *LacZ* reporter gene is used only to quantify the strength of the interaction, through quantification of β-galactosidase activity in cell extracts. To test the activation of the *ADE2* and *HIS3* reporter genes, diploid strains were plated on appropriate selective media without adenine or histidine, respectively, while to test the activation of the yeast *MEL1* reporter gene encoding a secreted α−galactosidase, haploid transformants (to check for autoactivation; Fig. S5) and diploid derivatives were plated on appropriate selective medium supplemented with X-α-Gal (Takara Bio) at a final concentration of 40 μg/ml.

To quantify activation of the *LacZ* reporter gene, a β-galactosidase activity assay was performed, using o-nitrophenyl β-D-galactopyranoside (ONPG) as a substrate. The assay was performed as described in the Yeast Protocols Handbook (Takara Bio). All reactions were performed in triplicate, so that they could be stopped at 3 different points in time (after 2 hours, 6 hours, and 24 hours), by the addition of 0.4 mL of 1M Na_2_CO_3_ to each suspension. Raw data concerning these assays is shown in **Table S8 (10**.**6084/m9**.**figshare**.**13176422)**.

### Construction of recombinant plasmids and gene deletion cassettes

For the construction of *P. rhodozyma* mutants, recombinant plasmids and gene deletion cassettes were constructed, as follows. Primer sequences (**Table S6 (10**.**6084/m9**.**figshare**.**13176422)**) were based on available genome sequences of *P. rhodozyma* strain CBS 6938 (NCBI project PRJEB6925 (58)) and *C. capitatum* strain PYCC 4530 (Bioproject PRJNA371774). Plasmids used for the constructions were pJET1.2/blunt (Thermo Scientific), pPR2TN containing a geneticin resistance cassette (59) and pBS-HYG (60) containing a hygromycin resistance cassette. All fragments used for cloning and deletion cassettes were amplified by PCR using Phusion High-Fidelity DNA polymerase (Thermo Scientific) and the amplified products were purified using either *Illustra GFX PCR DNA and Gel Band Purification Kit* (GE Healthcare) or *GeneJET Gel Extraction Kit* (Thermo Scientific). Constructions were performed using standard molecular cloning methods (61) and *E. coli* strain DH5α as host.

### Transformation of *P. rhodozyma*

*P. rhodozyma* strains (CBS 6938 and mutants) (**Table S4 (10**.**6084/m9**.**figshare**.**13176422)**) were transformed by electroporation, with the linearized recombinant plasmids or deletion cassettes, as previously described by Visser *et al*, 2005 (62), reducing the amount of DNA used to 2 μg for the transformations of the complemented *P. rhodozyma* mutants. The electroporation conditions (Gene Pulser II Electroporation System, Bio-Rad) consisted of an internal resistance of 1000 Ω, an electric pulse of 0.8 kV, a capacitance of 25 μF, resulting in a pulse ranging from 18 to 20 ms (63, 64). The cells recovered subsequently in YPD liquid medium, for at least 2.5 hours at 17 °C before being plated on YPD medium supplemented with the appropriate antifungal drug and incubated at 17°C. The genotypes of the transformants were determined as previously described (11, 65).

### *P. rhodozyma* sporulation (basidia formation) assays

To determine the ability of the *P. rhodozyma* mutants to sporulate, basidia formation assays were performed, where the strains were incubated in DWR solid medium with 0.5% of ribitol (0.5% Ribitol and 2.5% Agar), as previously described (11). Each basidia formation assay was conducted on 3 plates containing DWR+0.5% ribitol. On each plate, 10 colonies of each strain to be tested were spotted. Different strains were employed in each assay as indicated in **Table S2 (10**.**6084/m9**.**figshare**.**13176422)**, but in all assays the *P. rhodozyma* wild type was used as a positive control. Colonies were examined under the microscope using 100x magnification after 10 and 20 days of incubation at 18°C, and basidia formation patterns were scored qualitatively. The numbers of basidia counted in experiment E8 for the complemented mutants concerning the study of the *HD* locus of *C. capitatum* after 20 days of incubation are listed in **Table S3 (10**.**6084/m9**.**figshare**.**13176422)**. In all cases the entire colony was submitted to microscopic observation.

### Real-time quantitative PCR to assess expression of the *MAT* genes

Total RNA was extracted from a sporulating culture of *C. capitatum* strain PYCC 5626. Sporulation (teliospore formation in the case of this species) was induced by incubation in GSA liquid medium (0.2% Glucose, 0.2% Soytone) in 10% the volume of the flask, at 17°C, without agitation, until microscopic inspection revealed hyphae and teliospores, the latter being thick-walled resting spores from which basidia arise (16). Total RNA extraction was performed using the *ZR Fungal/Bacterial RNA MiniPrep kit* (by ZYMO Research). *P. rhodozyma* complemented mutants were grown in YPD liquid medium to an OD_600nm_ of 1.0. The cultures were then collected and frozen at −80°C for 1 hour before proceeding with total RNA extraction through a standard Trizol method. In-column DNase I digestion to free the RNA samples of genomic DNA using the RNA Clean & Concentrator kit (by ZYMO Research) was used for all samples, and absence of gDNA was verified by PCR. cDNA was synthesized from total mRNA using Maxima H Minus Reverse Transcriptase (Thermo Scientific) and oligo (dT)20 as primer. Real-Time PCR was performed using the SensiFAST SYBR No-ROX kit (by Bioline, London), with 20μl reactions, in a Rotor-Gene 6000 Corbett apparatus. The reaction parameters consisted of an initial denaturation step at 95°C, for 2 min, followed by 40 cycles of 95°C for 5 seconds, 57°C for 10 seconds and 72°C for 20 seconds. Two biological replicates were performed, with triplicates performed for each reaction (**Table S9 (10**.**6084/m9**.**figshare**.**13176422)**). Relative expression of the *MAT* genes was calculated using the 2^-ΔCt^ method, where ΔCt = Ct_test_ – Ct_reference_, as described by Livak and Schmittgen (66), and expression values were represented as the log_2_ (2^-ΔCt^). Mann-Whitney tests were performed to determine if the differences in expression of genes within each strain were statistically different.

## Data availability

Nucleotide sequences been deposited in NCBI/EMBL (GenBank) under accession numbers: MT561333-MT561342 (*C. capitatum STE3*); MT561330-MT561332 (*C. ferigula STE3*); MT592882-

MT592890 and MT592891-MT592894 (respectively for *C. capitatum* and *C. ferigula* 5’-end and intergenic region of *HD1* and *HD2*). Sequencing reads for *C. ferigula* PYCC 5628 (BioProject PRJNA371793) are available in the NCBI SRA database. The *C. ferigula* PYCC 5628 draft genome assembly has been deposited at DDBJ/ENA/GenBank under accession numbers MVAN00000000.

## ACKNOWLEDGEMENTS

The authors are very thankful to Alex Idnurm, Brenda Wingfield, Timothy James, and Paul Dyer for their insightful and thorough reviews of this manuscript. This work was supported by UCIBIO-Unidade de Ciências Biomoleculares Aplicadas, which is financed by Portuguese funds from Fundação para a Ciência e Tecnologia, Ministério da Ciência, Tecnologia e Ensino Superior (FCT/MCTES; https://www.fct.pt/) grant UID/Multi/04378/2019 and FCT/MCTES grants PTDC/BIA-GEN/112799/2009 (to P.G.), SFRH/BPD/79198/2011 (M.C.) and SFRH/BD/81895/2011 (M.D.P.). This work also benefited from the support of the INCD computing infrastructure funded by FCT and FEDER under the project 01/SAICT/2016 n° 022153. This work was also supported by NIH/NIAID R37 Award AI39115-23 and R01 award AI50113-16. J.H. is co-director and fellow for CIFAR program Fungal Kingdom: Threats & Opportunities.

## Supplementary Figures

**Fig. S1.**
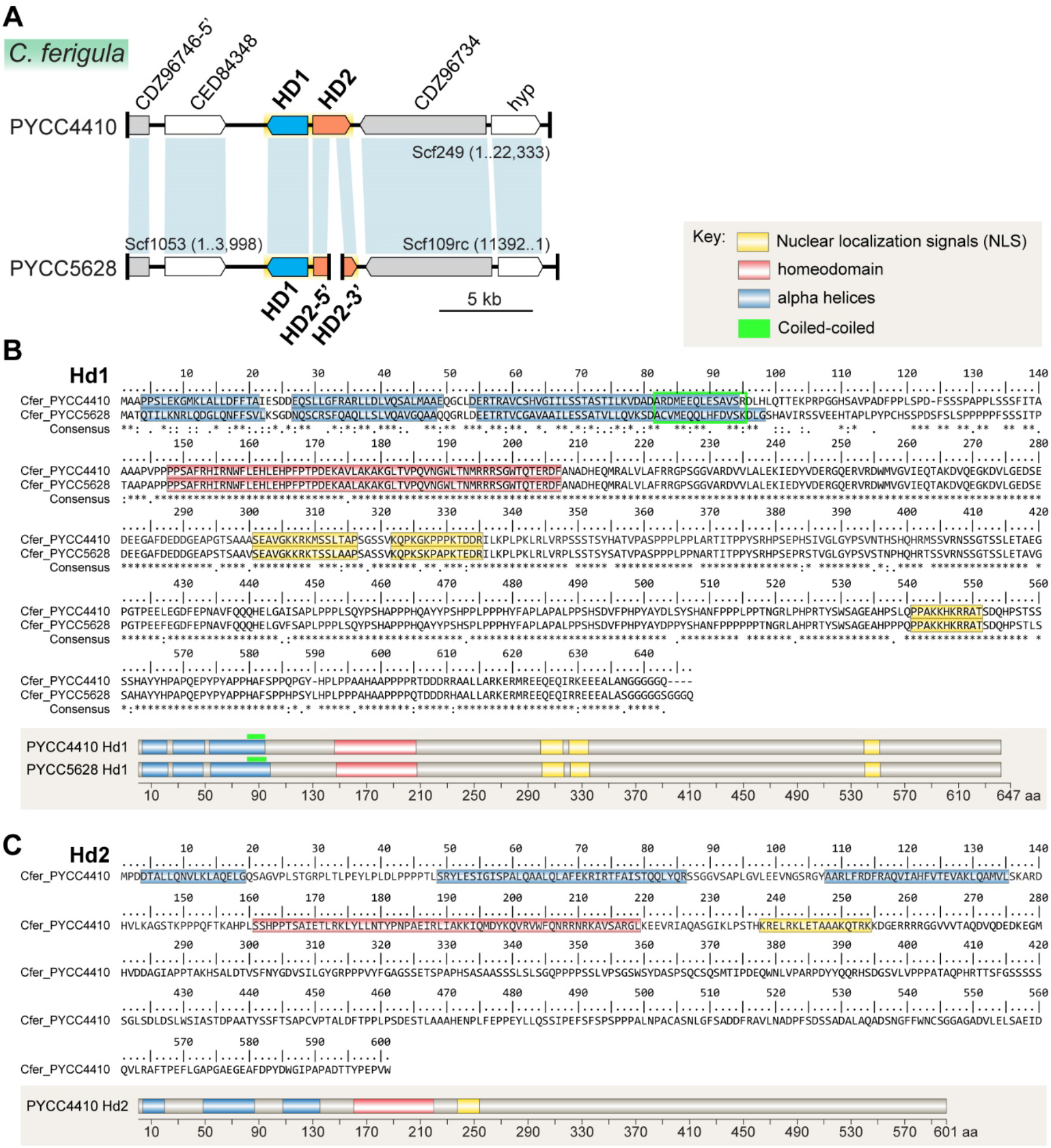
Analysis of the *HD* locus of *C. ferigula*. (A) Genomic organization of the *HD1* and *HD2* genes in *C. ferigula* PYCC 4410 and PYCC 5628. The *HD2* gene in strain PYCC 5628 is fragmented and localizes, as depicted, at the ends of two different scaffolds. However, comparative analysis indicates this region is syntenic in both *C. ferigula* strains. (B) Hd1 amino acid sequences of strains PYCC 4410 and PYCC 5628 shown as a sequence alignment (on the top) and as a schematic representation (on the bottom). (C) Hd2 amino acid sequence of *C. ferigula* PYCC 4410. In panels B and C, typical protein secondary structure features are highlighted according to the key on the top right.

**Fig. S2.**
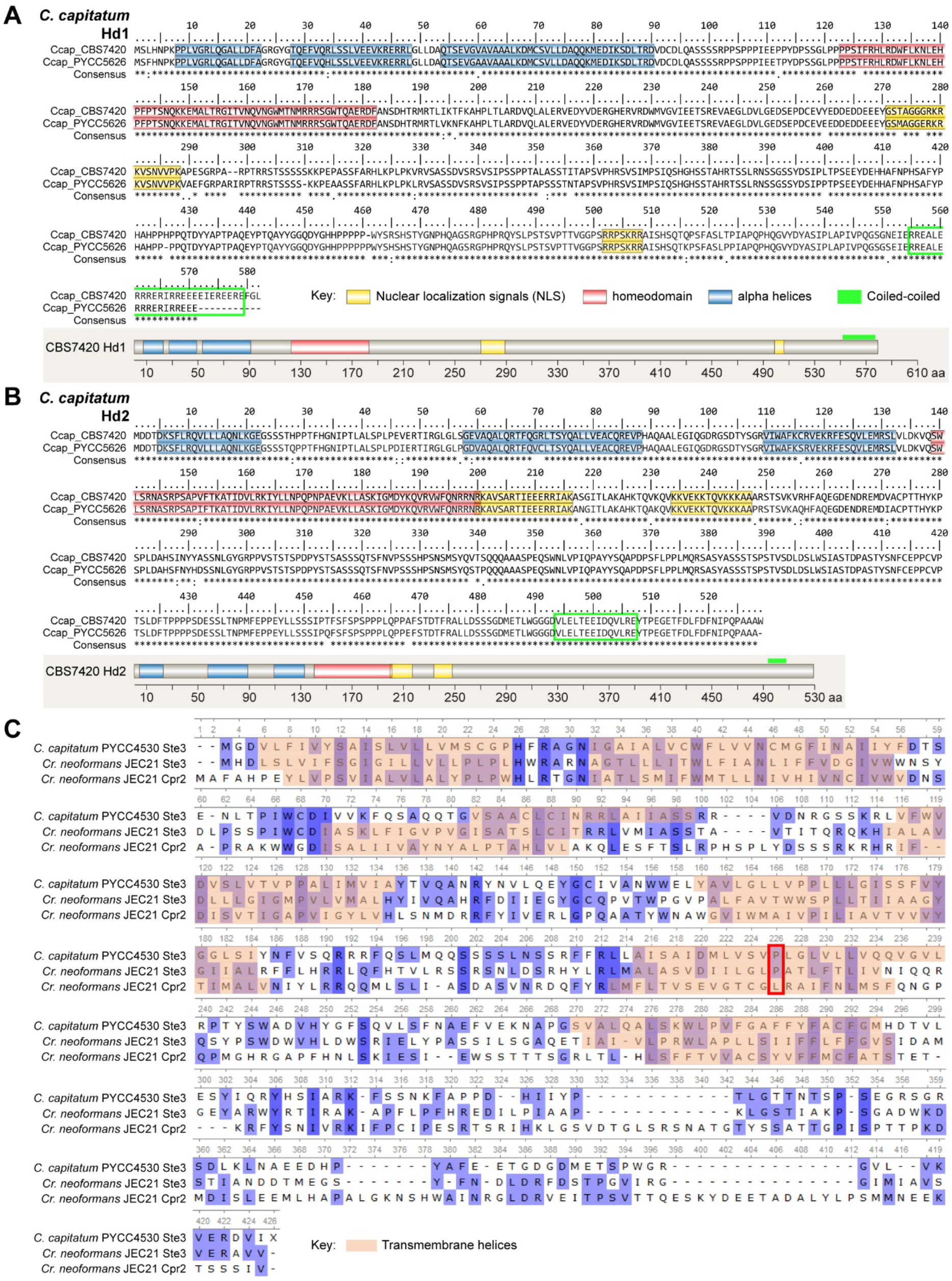
Sequence alignment of the (A) *HD1* and (B) *HD2* gene products from *C. capitatum* PYCC 4530 and PYCC 5626. In both panels, the amino acid sequence alignment is shown on the top and a schematic representation is shown below. Typical protein secondary structure features are highlighted according to the key on the bottom.(C) Sequence alignment of the Ste3 receptors from *C. capitatum* and *C. neoformans*, and the *C. deneoformans* pheromone receptor-like Cpr2, highlighting the seven transmembrane domains. The constitutive activity of Cpr2 in *C. deneoformans* results from an unconventional residue, Leu^222^, in place of a conserved proline in transmembrane domain six (red box; Hsueh Y-P, Xue C, and Heitman J, EMBO J 28:1220–1233, 2009).

**Fig. S3.**
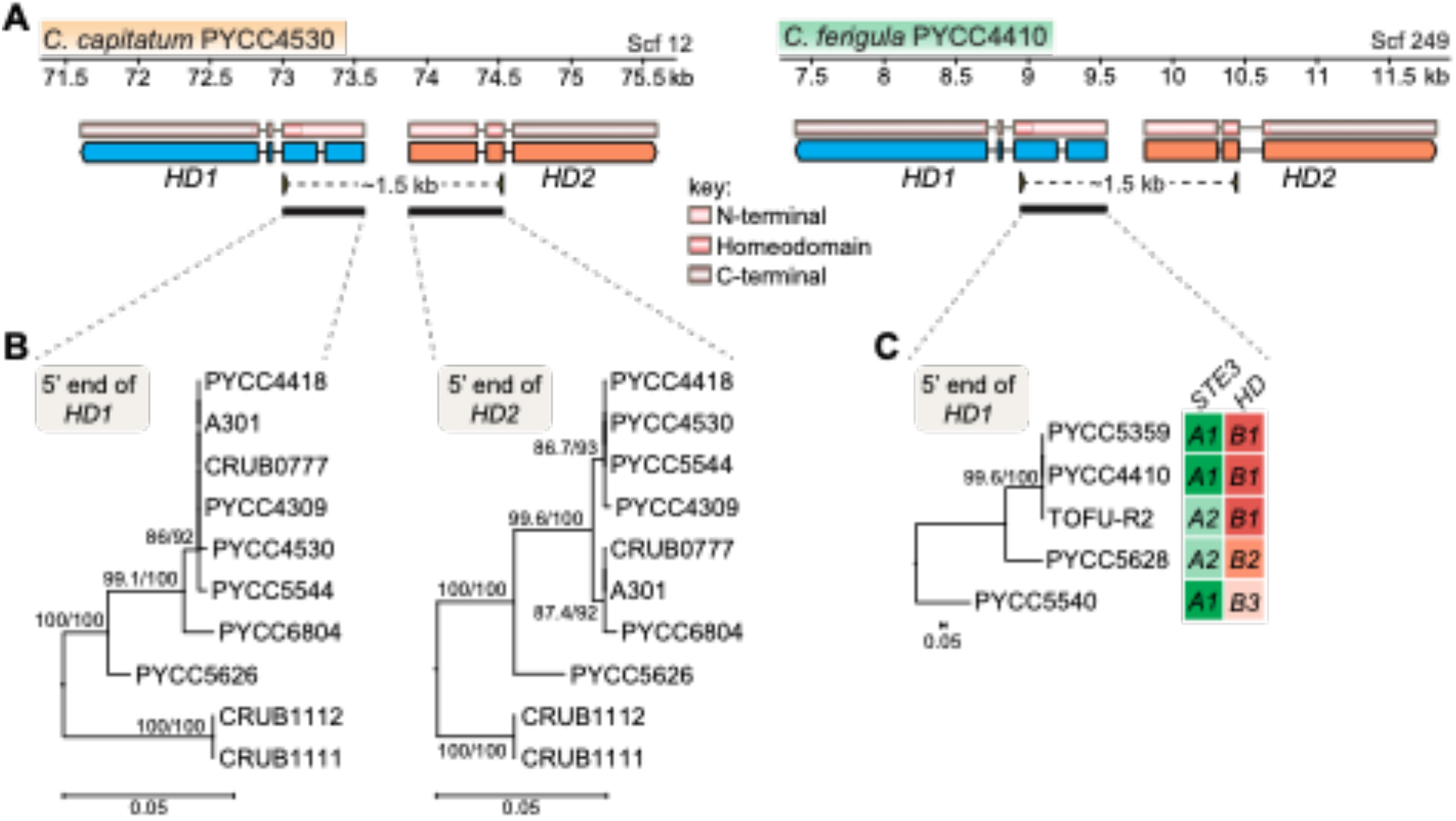
Determination of different alleles of the *HD* loci in *C. capitatum* and *C. ferigula*. (A) A ∼1.5-kb-long genomic region spanning the homeodomain, the 5’ end region, and the common intergenic region of the *HD1* and *HD2* genes was PCR-amplified and sequenced from available 10 strains of *C. capitatum* and 5 strains of *C. ferigula*. Primers locations are shown as yellow arrowheads below the genes. (B) Maximum likelihood phylogenies inferred from the 5’ end of the *HD1* and *HD2* genes (regions underlined in panel A) of *C. capitatum* using the best-fit substitution models HKY+F and K2P, respectively. (C) Maximum likelihood phylogeny inferred from the 5’ end of the *HD1* gene of *C. ferigula* using the best-fit model K2P. Note that the branch length in the two *HD1* gene trees is quite different (scale bars in nucleotide substitutions per site), which implies a much higher divergence of the *C. ferigula HD1* alleles compared to the *HD1* alleles of *C. capitatum*. The trees are rooted in the midpoint and branch support values separated by a slash were assessed by 10,000 replicates of both Shimodaira-Hasegawa approximate likelihood ratio test (SH-aLRT) and the ultrafast bootstrap approximation (UFBoot). The molecular mating type assigned for each of the analyzed strains of *C. ferigula* provides evidence that this species has a biallelic *P/R* locus and a multiallelic *HD* locus, which are genetically unlinked (the *B1* allele appears associated with both *A1* and *A2* alleles).

**Fig. S4.**
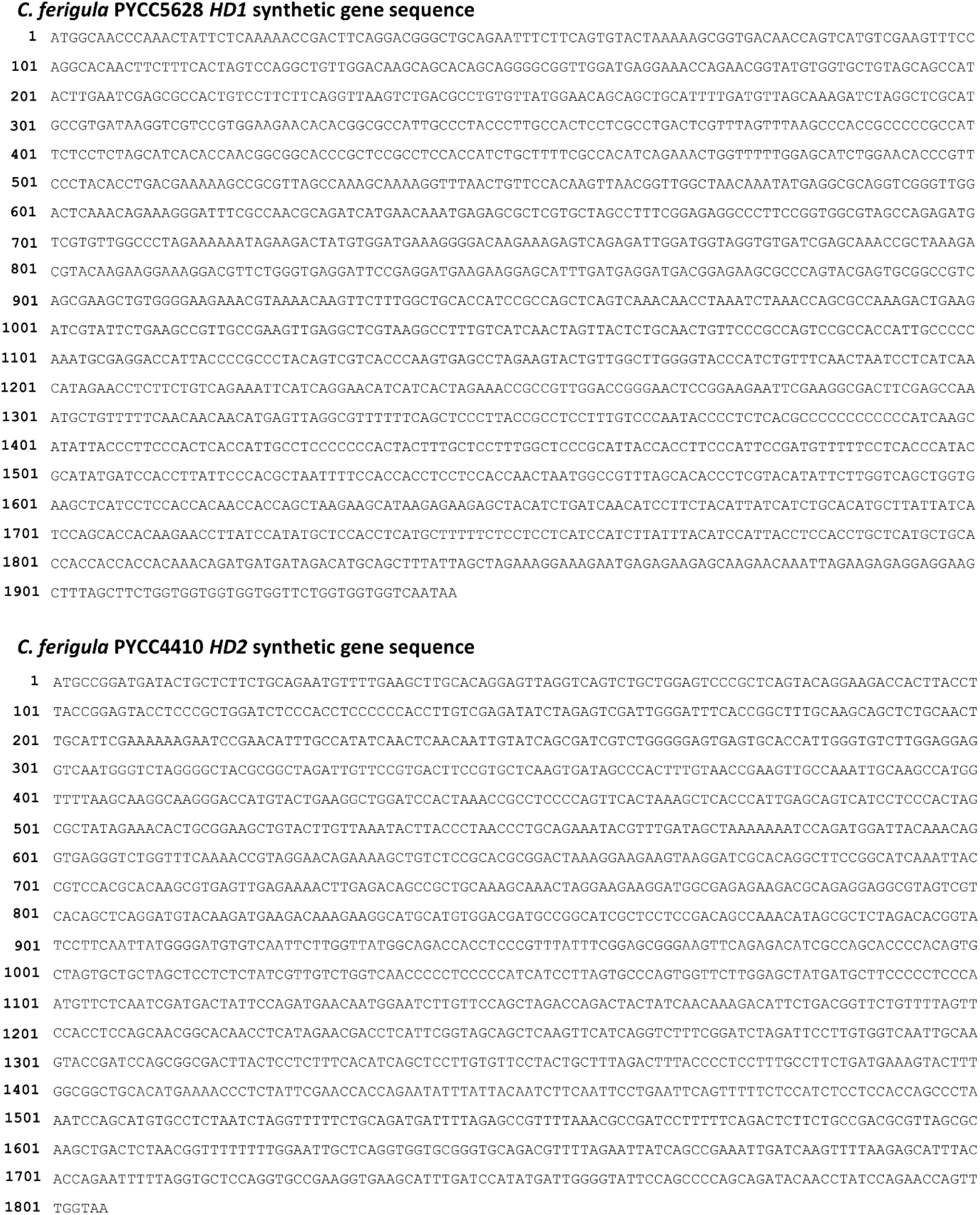
DNA sequences of the designed synthetic *HD1* and *HD2* genes of *C. ferigula* PYCC 5628 and PYCC 4410, respectively. Sequences correspond to the coding DNA sequences of the *HD1* and *HD2* genes of PYCC 5628 and PYCC 4410, respectively, optimized for the codon usage of *S. cerevisiae*.

**Fig. S5.**
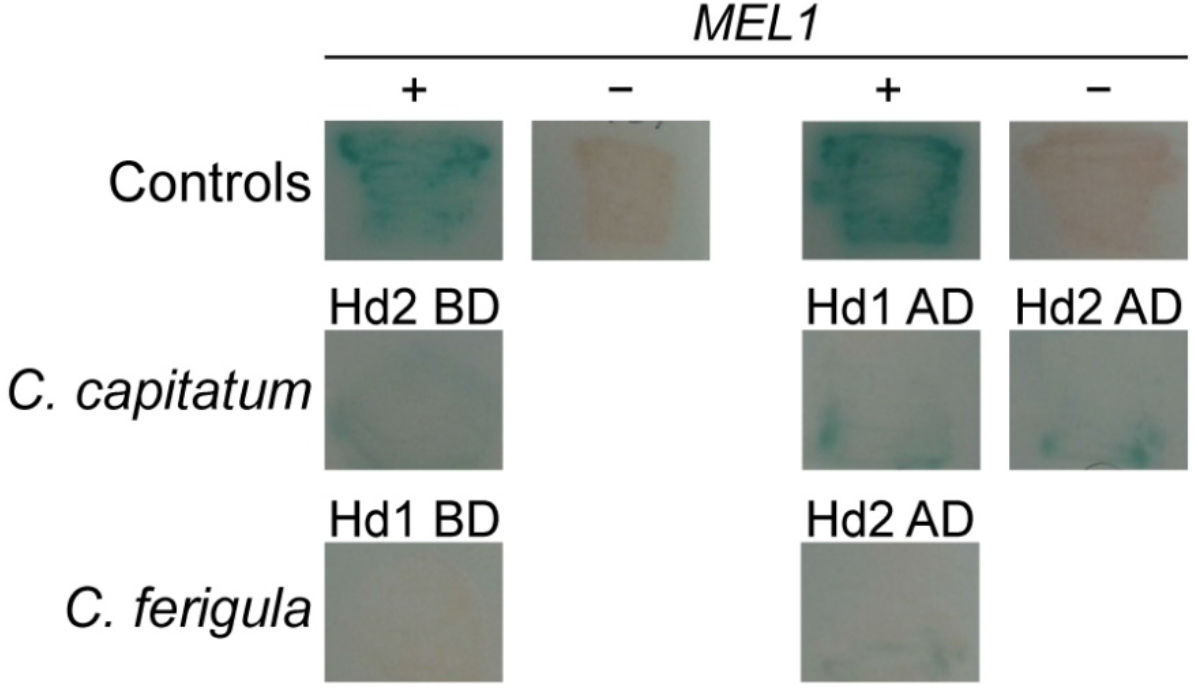
Activation of the reporter genes in the Yeast Two-Hybrid Assay by the individual fusion proteins. This control consists in assessing activation of reporter genes in haploid transformants carrying each of the fusion proteins to be tested. Absence of this so-called autoactivation shows that activation of reporter genes in diploid strains requires the presence of both interacting partners. Controls are the same as in Figure 4. AD, activation domain of Gal4; BD, DNA binding domain of Gal4.

